# Widespread reductions in body size are paired with stable assemblage biomass

**DOI:** 10.1101/2023.02.03.526822

**Authors:** Inês S. Martins, Franziska Schrodt, Shane A. Blowes, Amanda E. Bates, Anne D. Bjorkman, Viviana Brambilla, Juan Carvajal-Quintero, Cher F. Y. Chow, Gergana N. Daskalova, Kyle Edwards, Nico Eisenhauer, Richard Field, Ada Fontrodona-Eslava, Jonathan J Henn, Roel van Klink, Joshua S. Madin, Anne E. Magurran, Michael McWilliam, Faye Moyes, Brittany Pugh, Alban Sagouis, Isaac Trindade-Santos, Brian McGill, Jonathan M. Chase, Maria Dornelas

## Abstract

Biotic responses to global change include directional shifts in organismal traits. Body size, an integrative trait that determines demographic rates and ecosystem functions, is often thought to be shrinking in the Anthropocene. Here, we assess the prevalence of body size change in six taxon groups across 5,032 assemblage time-series spanning 1960-2020. Using the Price equation to partition this change into within-species body size versus compositional changes, we detect prevailing decreases in body size through time. Change in assemblage composition contributes more to body size changes than within-species trends, but both components show substantial variation in magnitude and direction. The biomass of assemblages remains remarkably stable as decreases in body size trade-off with increases in abundance.

**One-Sentence Summary:** Variable within-species and compositional trends combine into shrinking body size, abundance increases and stable biomass.

## Main Text

The loss or gain of large species can have dramatic consequences for ecosystem functions in terms of total system biomass and metabolism and thus food web energy fluxes (*1*). Anthropogenic changes to the biosphere are miniaturizing many communities (*1–3*) due to the extinction of larger species (e.g. (*4*)), and selective removal of the largest individuals (*5*). Yet shrinkage trends of community body sizes are by no means universal and, when looking across many communities, no change in body size is found on average (*6*). Indeed, as species shift their locations, some communities, including arctic and high-elevation plant assemblages (*7*), are also gaining relatively larger species. Hence, the prevalence of body size changes, and their implications for assemblage abundance and biomass are unknown.

Here, we first quantified the distribution of trends in body size change through time from 5,032 time-series over 60 years (*8*), including data from 4,734 species and six taxon groups from communities distributed across the world (Fig.1A, fig. S1). We find more shrinking than increasing trends, both overall, and among time series with stronger evidence for trends of change (lower p-values) (Fig. 1B), contrasting with the results of Terry et al. (*6*). To disentangle the variety of trends in body size we found and how they align with previous studies, we decompose body size changes into the different mechanisms involved, namely the loss or gain of species of different sizes (i.e., compositional change) and within-species population changes as individuals of a given species tend towards smaller or larger sizes.

**Fig. 1.**
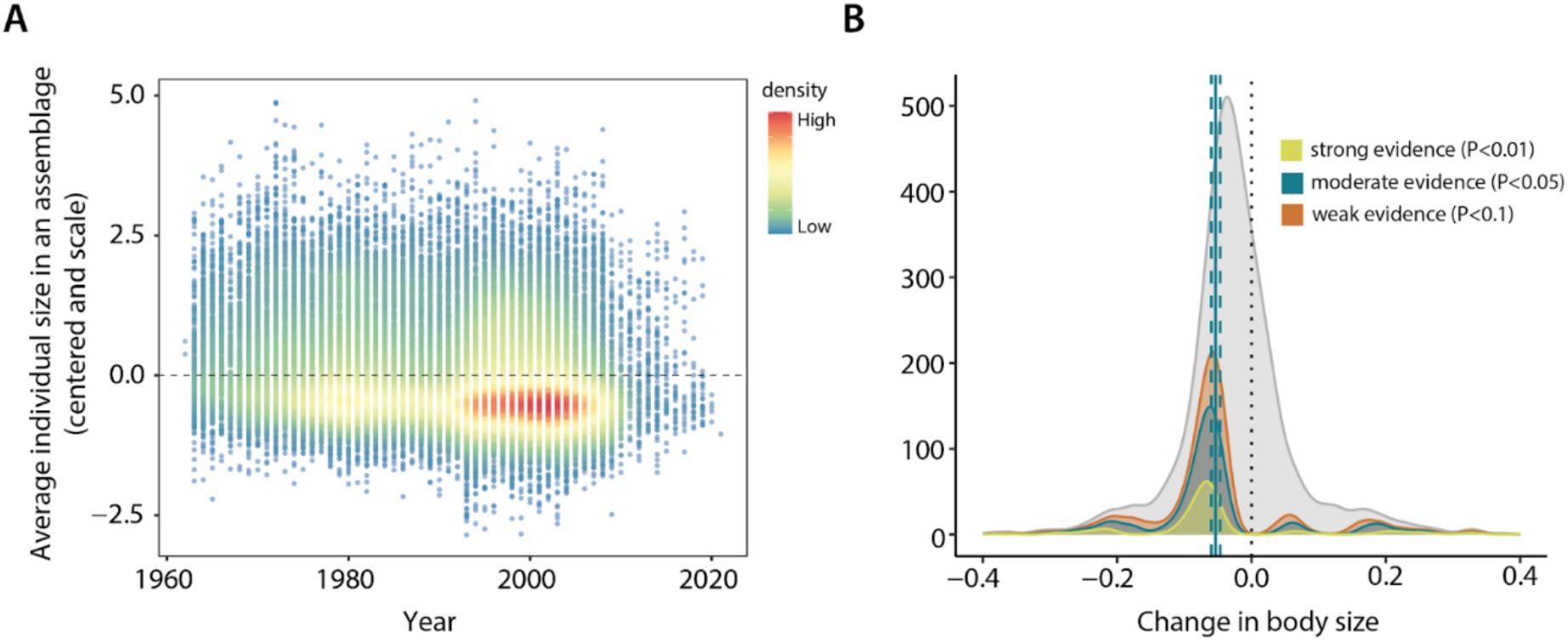
Changes in body size across 5,032 assemblages reveal higher prevalence of declining trends. **(A)** Average individual size across the full set of assemblage time-series. Each point represents an assemblage body size at one time point, coloured by density break (colder colours indicate lower densities) **(B)** Density plots of the distribution of slopes of change in average individual body size. The full set of 5,032 assemblage time-series is shown in light gray. Yellow, blue and orange represent respectively the subset of assemblages for which strong evidence (P<0.01), moderate evidence (P<0.05) and weak evidence (P<0.1) of change was detected when testing slopes against 0. Dotted lines show slope of 0, while blue dashed lines show the mean slope across the blue data (traditional significance value) and the respective 90% credible interval.

Population body size shifts have been attributed to several selection forces, including preferential exploitation of larger individuals (*1, 9*), climate change and habitat conversion (*10–12*) At the assemblage level, large-bodied species have been particularly susceptible to extinction following temperature shifts and human colonisation of landmasses due largely to their life history traits and lower numbers (*13*), which can shift composition turnover to favour small-bodied species. Previous assessments often investigate only compositional (*6*)components of body size change or within-species population changes (*3*), but see (*14*). We collated time-series that recorded both organism abundance and body size data in the field across 5,032 assemblages to estimate the extent to which the two mechanisms drive overall change in body size across taxa and regions.

Body size change from compositional and within-species change can co-occur, and operate either in the same or opposing directions (Fig. 2). We decomposed assemblage body size change into compositional and within-species changes using an extension of the Price equation (*15–17*). The Price equation is a mathematical description of the relationship between statistical descriptors (mean and covariance) of selection and trait change (*18*). By examining the type of covariance in these two mechanisms of change, we can determine the relative contributions of compositional and within-species change to observed overall change. Assemblage body size change is most pronounced when both mechanisms operate in the same direction (towards either shrinking or increasing body size), such that the covariance between compositional and within-species changes is positive. When only one mechanism is involved (i.e., change in one axis but not the other), body size change tends to be lower; and with negative covariance it is possible to have change in one component cancel out change in the other. (Fig. 2).

**Fig. 2.**
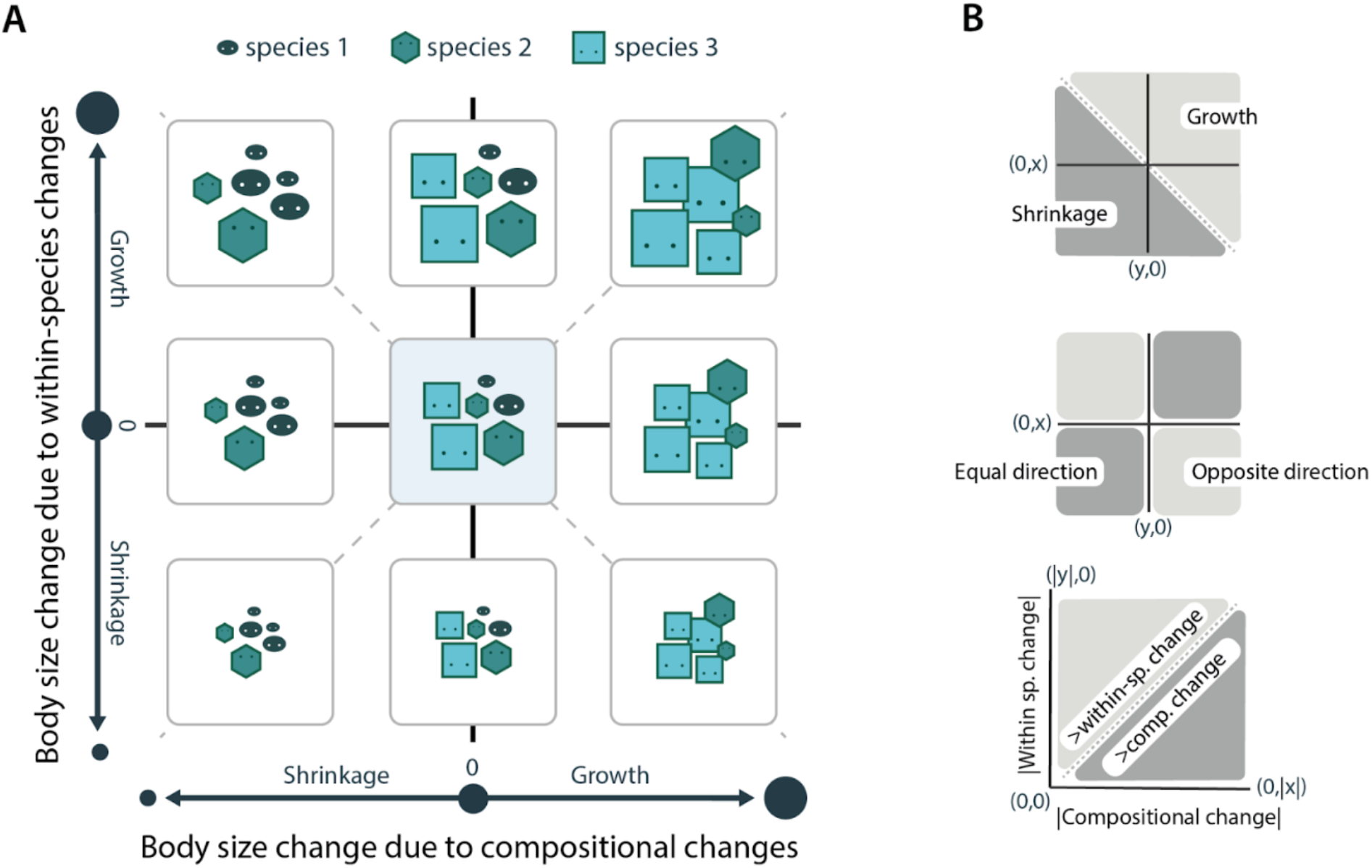
Mechanisms that underpin temporal changes in mean body size. **(A)** Shifts in mean assemblage-level body size can occur due to within-species changes (vertical axis), compositional changes (horizontal axis), or a combination of both mechanisms, displayed as change between two time points: from time 1 to time 2. Body size distribution for time 1 is shown in the middle, with different change outcomes for time 2 shown in the other cartoons, omitting the no-change scenario where time 2 is identical to time 1. Different colours and shapes represent different species. Icon size represents the body size of an individual. Note that: The vertical placement (axis) represents the within-species (population or intraspecific) changes through time in mean body size. This could be a mix of increases (e.g., smaller individuals growing larger or being replaced by larger individuals) and decreases in the average size of individuals of different species. The horizontal placement (axis) indicates change in mean body size resulting from the gain or loss of species (compositional turnover), or a change in the relative abundance of the species present in an assemblage (even without local extinction or immigration of species). **(B)** Changes are not expected to follow a generalised directional pattern; the two components can reinforce or counteract each other. If they counteract, the overall direction of change will depend on which component shows the higher absolute effect (contribution).

Our analysis shows that assemblage body size is predominately shrinking, with substantial variation in the balance of within-species change and compositional change (Fig. 3). Of the 5,032 assemblages, two-thirds decreased in assemblage average body size and one-third increased. In fact, both mechanisms of body size change were present for the overwhelming majority of assemblages (97%) with the magnitude of compositional change being greater than within-species change in 72% of assemblages. Although compositional and within-species change often occurred in the same direction (59% of assemblages), we found counteracting effects in 41% of all assemblages. For example, of the 3,478 assemblages showing within-species decreases in body size, compositional change for ~37% drove increases in body size (when within-species change < 0 and compositional change > 0, Fig 3). Consequently, the conflicting trends in body size previously reported in the literature are likely linked to this substantial variation in magnitude and direction of the two components of body size change.

**Fig. 3.**
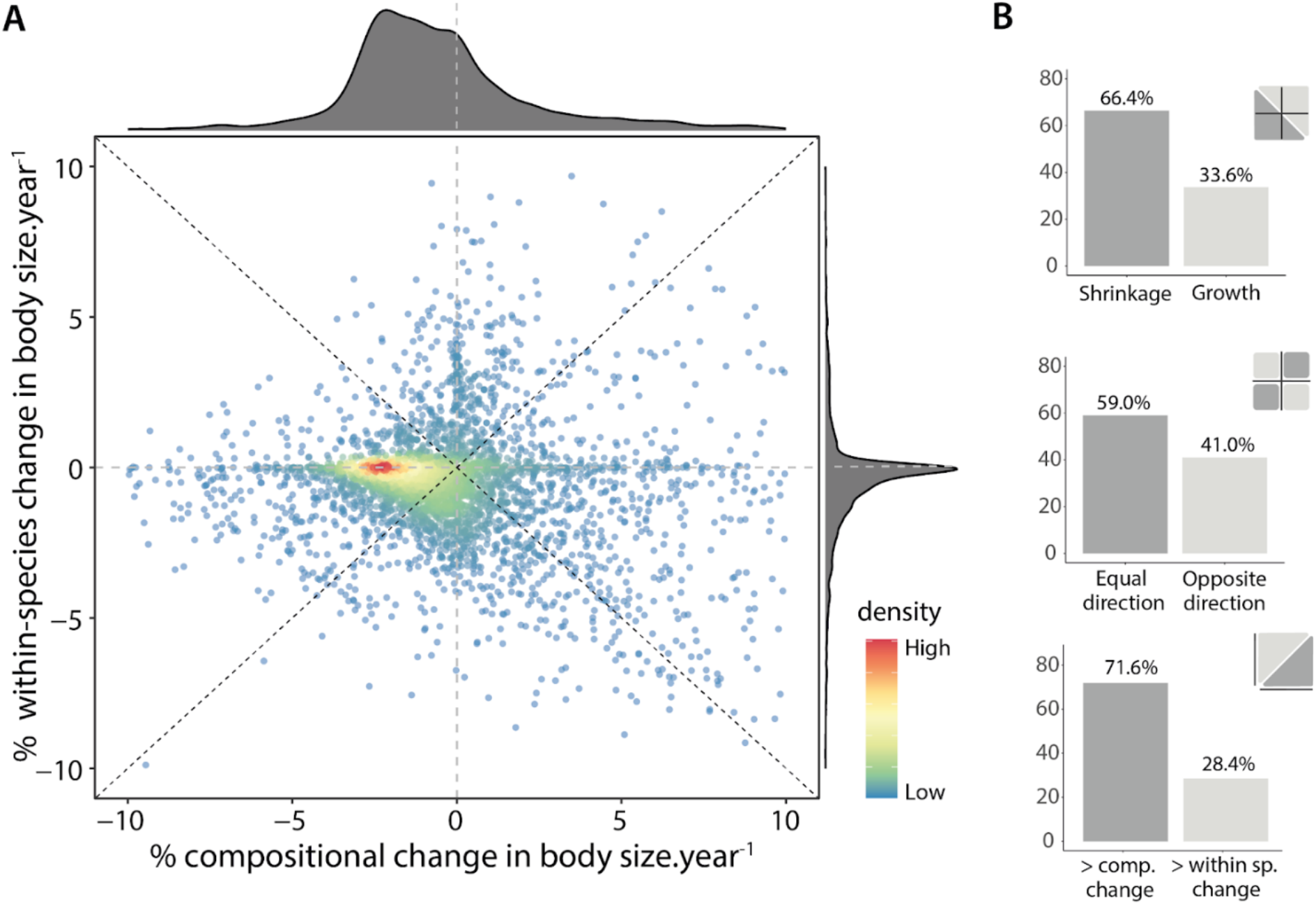
Patterns of body size change through time in 5,032 assemblages representing 4,237 species of fish, plants, invertebrates, mammals, herpetofauna, and marine benthic organisms. **(A)** Relationship between population-level (i.e., within-species) changes and assemblage-level (i.e., compositional) changes. Both axes show % changes standardised by the number of years between the first and last year sample of the assemblage (duration); assemblages (points) are coloured by density break (colder colours indicating lower densities). Dashed lines show x = 0, y = 0, x = y and y=-x. **(B)** Frequency distributions (in percentage) of the number of assemblages (n=5032) in the different scenarios depicted in Fig. 1B. For clarity, assemblages with % change.year^-1^ higher than 10% (n=514; but see fig.S2 & S5) are not shown in panel (A), but are included in (B).

We found that trends in body size change over time differ according to taxa, realm and latitude (fig. S3). Available data with simultaneous estimates of abundance and body size displays geographic and taxonomic biases (fig. S1), with order of magnitude differences in the number of species, observations and time-series among different taxa. Hence our confidence in estimates of body size change is highest for the most well sampled taxon—marine fish, which show a particularly evident decrease in body size (fig. S3A). This result aligns with previous evidence of directional trait changes found among fish assemblages (*19–21*). For marine organisms, these changes are often linked to the selective exploitation of large-bodied individuals by humans (*19*), to warming (*20*), or to decreased resource availability (*21*). Disturbances and selective removals affect the age structure of populations, as well as genetic shifts within populations (e.g., (*22*)). Combinations of these drivers likely result in high variability in trends and prevalence of counteracting effects. Among other taxa, realms and climates (fig. S3) the number of available time-series is lower and within-species and compositional changes are more variable. We see evidence of increasing body size where the pattern of compositional change is strongest (e.g., plants, invertebrates and benthos-mainly marine invertebrates), counteracting declining trends within species, although these results should be interpreted with caution given the lower number of time-series involved. There is substantial variability in the duration and time period of the time-series in our dataset, but our results are robust to the length of the time-series as well as the start and end points (see sensitivity analysis in fig. S4).

Body size is usually tightly linked to abundance (*23*) through metabolic (*24*) and trophic (*25, 26*) processes. This relationship can have implications for assemblage biomass mediated by a trade-off between size and abundance (*27*). Hence, we investigated if the changes in body size were associated with changes in assemblage abundance, biomass, or both. We found that abundance has, on average, slightly increased through time, while the overall change in biomass is indistinguishable from zero (Fig. 4). Previously, no (*6*), or complex (*28*) relationships have been found between body size and abundance changes. Our results also point to a complex relationship. However, amidst the variation, there are signs that the overall reduction in body size is being counteracted by increasing overall abundance (Fig. 4A and C). Trade-offs between abundance and body size are expected (*23*) and affect ecosystem metabolic rate and function (*24*). We detect a strong positive covariance between change in biomass and abundance (Fig. 4D), but much weaker covariance between abundance and body size, with the strongest trends among these two variables tending to negative covariance (Fig. 4C). In fact, 76% of the assemblages with detectable trends in both variables have abundance increases and body size decreases. These patterns suggest that assemblage body size, abundance and biomass are linked so that change in one has implications for change in the others. Evidence of widespread regulation of assemblage level variables (species richness and abundance) has been previously reported (*29*), whereby assemblages tend to return to previous levels after disturbances. The absence of a directional trend in biomass suggests it may be more tightly regulated than body size and abundance, which may be causing tradeoffs in change of the latter two variables.

**Fig. 4.**
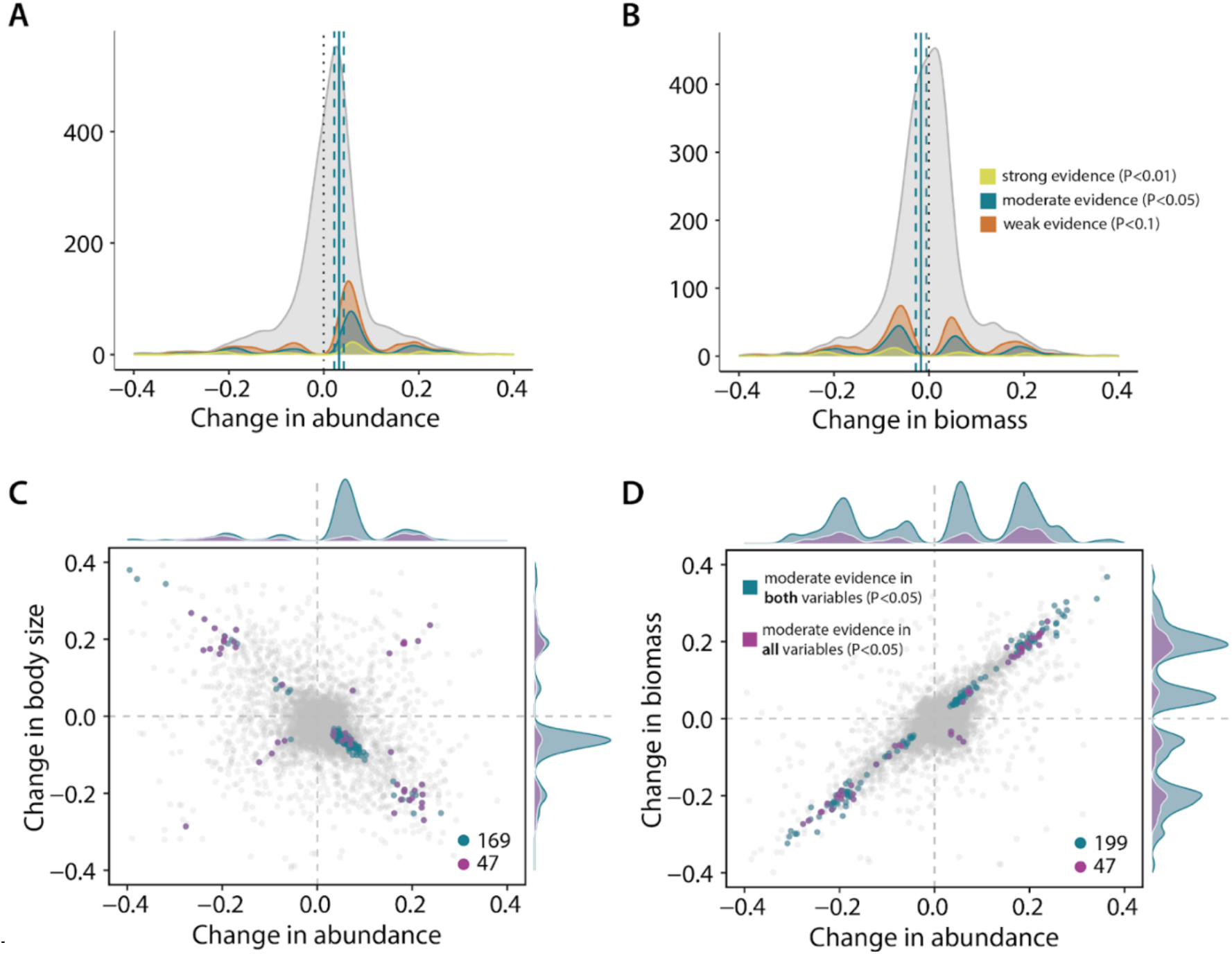
Changes in assemblage abundance, biomass and body size. Density plots of the distribution of slopes of **(A)** change in total abundance of individuals (of all species), and **(B)** change in total biomass in an assemblage, as a function of time, for the same assemblages as shown in Fig. 3. The full set of 5,032 assemblage time-series is shown in light gray. Yellow, blue and orange represent respectively the subset of assemblages for which strong evidence (P<0.01), moderate evidence (P<0.05) and weak evidence (P<0.1) of change was detected when testing slopes against 0. Dotted lines show slope of 0, while blue dashed lines show the mean slope across the blue data (traditional significance value) and the respective 90% credible interval; **(C** and **D)** The bottom panels show the different relationships between variables. Only assemblages for which strong or moderate evidence (P<0.05) were detected for both variables plotted are shown in blue, while purple highlights the assemblages for which significant changes through time were detected in all 3 variables (n=47), all remaining assemblages are shown in light grey: **(C)** change in average body size as a function of abundance changes (note that 76% of the blue dots are in the quadrant where abundance increases and body size decreases), and **(D)** change in biomass as a function of abundance changes.

Collectively, these analyses reveal high variation in local-scale assemblage body size change patterns, with both within-species and compositional changes playing considerable roles. These findings highlight the importance of considering interactions between compositional and within-species body size change. Specifically, the assemblage context is needed to understand within-species change. For example, removing top predators (often the larger-body size individuals in an assemblage) can trigger mesopredator release, which alters assemblage size structure and composition (*30*). It is also at the assemblage level where regulation will play out, for example, in relation to ecological carrying capacity (*31*). Similarly, compositional change on its own ignores important within-species variation. In our analyses, although we detect shrinkage with *in situ* estimates of body size, we cannot detect change when we use species’ mean body sizes (from external non *in-situ* trait databases) instead (fig. S8-S10) (*17*). Indeed, we detect no trend regardless of whether we use the same subset of the BioTIME database as in our main analysis, or if we expand it to include 20,925 assemblages (fig. S8) as was also found by (*6*). This discrepancy cautions against the use of species’ averages for traits that exhibit substantial individual variation (*20, 32*) and highlights the need for increased surveying of individual-level traits. Importantly, considering the two axes of variation (compositional and within-species) avoids the misleading conclusions that arise when the two components change in opposite directions.

The selection forces acting on body size are varied and have heterogeneous distributions in space and time. Partitioning body size change into within-species and compositional change, the two components adopted here, can help explain the wide variation in body size change patterns found in literature. For example, global warming is simultaneously selecting for smaller body size (for metabolic reasons), affecting species’ phenology, and causing range shifts (*2*). Global warming and species phenology effects can best be seen in within-species changes while range shifts induce compositional change. The net result of these processes will depend on the environmental context: in the Arctic Tundra, warming promotes larger shrubs (*7*), because southern species are expanding their ranges and because there are longer growing seasons. In contrast, warming is associated with smaller fish in the North Sea (*11*), although selective harvesting/exploitation is likely also contributing to this change (*33*). By considering both within-species and compositional changes in individual-level body size, alongside changes in relative abundance, future research should be able to better elucidate the mechanisms involved in body size change.

In conclusion, we find evidence of widespread shrinking body size through time due to various mechanisms despite substantial variation, and overall stable assemblage biomass. Body size is an easily measured, integrative, key morphological trait that scales with many ecological characteristics of organisms and ecosystems, such as demographic rates, metabolism and resource requirements (*34, 35*). We reiterate pleas for more regular monitoring of body size (*36*), especially for non-marine taxa. Future research should focus on the implications of the changes we identify for ecosystem function. For instance, cascading food web effects of shrinking body size in organisms could negatively affect human nutrition and associated economics (e.g., affecting crop plants and protein sources such as fish; (*37*). Moreover, shrinking body size through compositional change is likely to bring changes in other traits, and therefore trigger additional impacts on ecosystem functioning (*12*). Our study suggests the ubiquitous turnover in composition currently unfolding (*38, 39*) is a profound reshuffling of not only species, but also key characteristics of living organisms.

## Methods

Our analysis is based on bringing together trait data and ecological assemblage timeseries. We used two sources of body size trait data: direct measurements of species’ body size (biomass) taken over time in the field (type 1) and species’ average body size estimates from major published trait databases (type 2). Below, we describe the *a priori* quality criteria, standardisations and subsequent calculations and statistical analyses that we used to provide a uniquely comprehensive analysis of the processes behind current assemblage- and population-level body size changes using type 1 data. The results of these analyses are presented in the main text. In addition, we also present the methodology we followed to explore the global patterns of body size change when considering compositional changes alone and using different types of trait data (type 1 and type 2); the main results of those analyses are presented in the Supplementary Results (*17*).

### Assemblage time-series data (BioTIME Database)

BioTIME is the largest global, open-access database of assemblage composition timeseries (*8*). This database includes data on multiple multicellular taxa (e.g., plants, fish, birds, mammals, invertebrates), with over 12.5 million species-level records representing ~46,000 species. Each BioTIME study contains distinct samples measured (with a consistent methodology) over time, which could be fixed plots (i.e., ‘single-site’ studies where measures are taken from a set of specific georeferenced sites at any given time) or wide-ranging surveys, transects, tows, and so on (i.e., ‘multi-site’ studies where measures are taken from multiple sites that may or may not align from year to year). Because the spatial extent varies across studies, we followed previous approaches (*38, 40*) to identify and standardise ‘multisite’ studies using a global grid of 96km^2^ hexagonal cells using dggridR (*41*). Studies that were contained within a single cell were not partitioned. Following this step, each sample was assigned a different combination of study ID and grid cell (based on its latitude and longitude) resulting in a unique identifier for each assemblage time-series within grid cells, thus allowing for the integrity of each study and each sample to be maintained. Then sample-based rarefaction was applied to standardise the number of samples per year within each time-series (*38, 40, 42*). Finally, we only retained observations sampled after the year 1960 (99.4% of all the data was recorded after this period) and restricted our analysis to time-series with at least five different sampled years. For the analyses presented in the main text, we further subsetted the database, and considered only records that contain both abundance information (i.e., counts of the number of individuals) and biomass estimates. In total, we considered 5,032 time series from 44 studies across the globe (fig. S1; Table S1).

### Trait data (BioTIME Database)

We extracted body size trait data directly from BioTIME using both abundance and biomass estimates measured at the same time and place. From each record *i*, we estimate average body size of individuals (BS) by considering that:

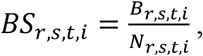

where *B* is biomass and *N* is abundance recorded in year *t*, for species *s* within the assemblage time-series *r*. Note that biomass is only measured at the individual scale (abundance=1) for ~28% of the data, thus we refer to this measure as average individual body size. Here, we include all taxonomic groups for which the appropriate data were available including groups poorly represented in most trait compilations (e.g., invertebrates). In total, we considered the following taxon groups: fish, benthos, plants, mammals, invertebrates, and herpetofauna (reptiles and amphibians). We did not distinguish taxa across realms, but studies that included multiple taxa (i.e., studies that sampled multiple taxon groups simultaneously) were re-classified based on the dominant taxa represented. On average, we estimated multiple body size measurements for 4,734 species within the 5,032 assemblage time-series (average ~41 species per assemblage time-series; Fig. S1).

### Partitioning body size change – Price equation

To partition temporal changes in average individual body size, we used an extension of the Price equation (*15, 18*), that allows an exact partition of trait change, Δz, in an observed dataset:

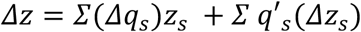

The first term on the left-hand side of the equation for Δz accounts for total body size change caused by changes in frequency due to species selection, i.e. changes in the relative abundance or presence of species with a certain property value (body size; e.g., local extirpations or colonisations). This term reflects the effect of species turnover. The second term describes the part of total change caused by changes in mean property values, reflecting the effect of within-species variation (e.g., larger individuals within a species being replaced by smaller individuals of the same species). Together, the two terms sum up to the actual change in community-weighted mean (CWM) body size in an assemblage, Δz. Given this, we quantified changes in frequency as *Δq_s_=q’_s_*-*q_s_*, where *q_s_* and *q’_s_* are, respectively, the before and after relative abundance of species *s*; and changes in mean property value as *Δz_s_*=*z’_s_*-*z_s_*, where *z_s_* and *z’_s_* represent, respectively, the before and after mean individual body size of species *s*. In assemblage time-series where multiple individual *BS_r,s,t,i_* estimates were available for the same year and species (e.g. when the assemblage was monitored more than once a year), an abundance-weighted mean was used instead. Finally, when *z_s_* was not available (i.e. colonisations did not occur) we considered *z_s_*=*z’_s_* for that species, and thus *Δz_s_* is equal to 0 and no change occurs due to changes in mean property values (within-species changes).

For each assemblage time-series, we used the Price equation to partition body size changes that occurred between two years, the last year (*t*_2_) and the first year (*t*_1_), where *t* = *t*_2_ – *t*_1_ + 1 is the full length of the assemblage time-series. In order to ensure comparability among time-series of different durations, both the within-species and the composition component of body size change were converted to proportional changes relative to the starting size of the assemblage. This was done by dividing each component of change by the initial assemblage CWM and standardising it by duration (i.e., dividing by t). These quantities were expressed in units of % change.year^-1^. Patterns across all assemblages are represented in Fig. 3 and fig. S2, and patterns across the different taxa, realms and climates are shown in fig. S3.

### Sensitivity analyses

Many of the assemblage time-series varied in length (27.4 ± 12yr, mean ± sd), with varying start and end points. To examine whether our results were sensitive to such effects, we repeated the analysis using alternative start (t_1_) and end times (t_2_) within the same assemblage. This analysis included a scenario where the first year was fixed, a scenario where the last year was fixed and a scenario where both years varied randomly. For each of the three scenarios, we repeated the analysis 100 times, where for each iteration we used the Price equation to partition body size changes that occurred between the selected two years in a given assemblage (as done in our main analysis); and reported the median effect of each component and their dispersion (interquartile range) across all iterations (fig. S4). Despite slight differences across scenarios, the results were largely concordant and yielded the same directional trends.

Given that some of the estimates were extreme with very large changes to assemblage level body size (fig. S2), there could be some concerns about errors in the measurement or measurement reporting of the abundance and biomass estimates (and consequently body size estimates) in the original datasets. We performed in-depth checks of the raw data within affected individual datasets (see fig S5 for an example) and found that such effects seem to be a true representation of changes occurring in the assemblages. Nevertheless, these few assemblages have the potential to over-influence the overall effects found, so we chose to report robust statistics (median and interquartile range) that de-emphasise such extreme cases without removing them, when appropriate (e.g., fig. S4 and S6).

For a few assemblage time-series the sample-based rarefaction process could lead to a different species composition. To ensure our results were robust to the random samples selected by the sample-based rarefaction process, we performed a bootstrap analysis rerunning the analysis described in the main text (using first and last year only) 100 times, each time using a different dataset after the sample-based rarefaction process was applied. Only the results of one iteration are presented in the main text, but plots of the distribution of results across the 100 rarefaction iterations can be seen in fig. S6.

### Body size, abundance, and biomass change

As mean body size emerges from the ratio of biomass and abundance (see section *“Trait data (BioTIME Database)”*), change in either biomass or abundance can be responsible for any observed body size changes. To explore these effects, we quantified trends in biomass and abundance across individual assemblage time-series. This was achieved by fitting ordinary least-squares regression models for each assemblage separately, with either average individual body size, total abundance, or total biomass (centered and scaled) as a function of time (year, mean-centered). All sampled years were considered, and for each year in a given assemblage time-series, total abundance and total biomass were calculated by tallying the number of individuals and biomass (regardless of species) sampled within that year, respectively. The average individual size in a year was then retrieved by dividing the sum of the biomass by the total abundance reported in that year for that assemblage time-series. The set of slopes (β) of these linear models is shown in Fig. 1 (change in body size) and Fig.4A and B (change in abundance and biomass, respectively). Additionally, we evaluated the associations between the temporal trends in total abundance, total biomass, and mean body size, by comparing the slopes of change of assemblages for which statistically significant trends were found across two or more variables (Fig. 4C and D; fig. S7). All calculations and statistical analyses were performed in R-3.6.3 (*43*).

## Author contributions

Conceptualization: all coauthors; Data curation: **I.S.M., V.B., C.F.Y.C., R.vK, A.B., A.F-E. and F.M.;**Methodology: all coauthors; Formal analysis: **I.S.M.;**Supervision/Project administration: **I.S.M.**, **M.D., F.S., and J.M.C.;**Visualization: **I.S.M.**, **M.D., F.S., C.F.Y.C., B.P., A.F-E., A.E.B., M.M., and S.A.B.;**Writing – original draft: **I.S.M.**, **M.D., F.S., R.F. and J.M.C;**Writing – review and editing: all coauthors;

## Data and materials availability

The BioTIME data can be accessed on Zenodo (https://doi.org/10.5281/zenodo.2602708) or through the BioTIME website (http://biotime.standrews.ac.uk/); The selected data used in this article and the R scripts used to generate the main results of the study are available through (https://doi.org/10.5281/zenodo.7595981).

## Supplementary Materials

Figs. S1 to S10

Supplementary Results

Table S1

References (*44–258*)

