## Supplementary Materials for "Widespread reductions in body size are paired with stable assemblage biomass"

**This PDF file includes:**

Figs. S1 to S10
Supplementary Results
Tables S1
References (*44–258*)

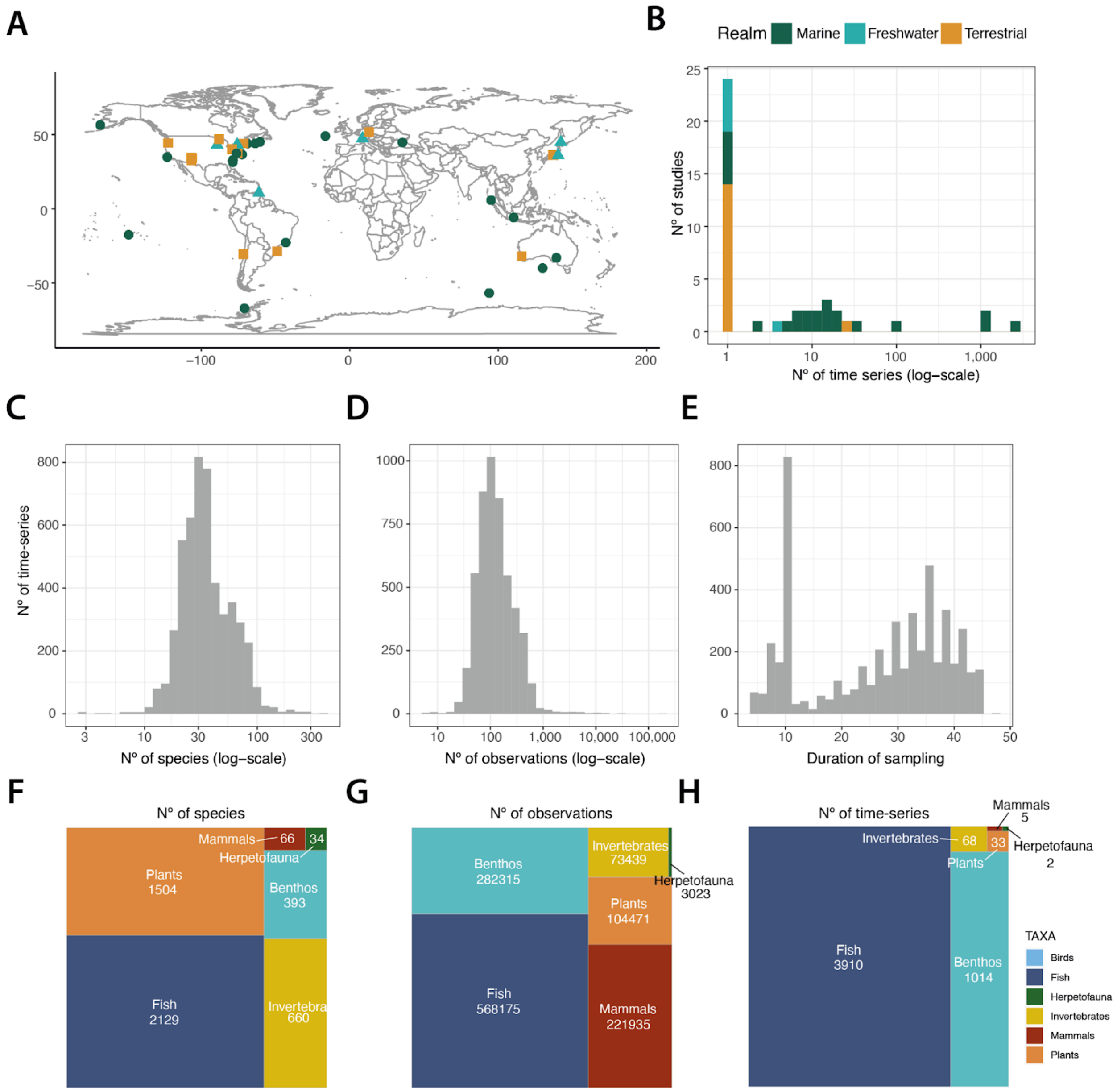


**Fig.S1.**

**Distribution of study data characteristics (BioTIME data, type 1). (A)** Location of the studies with direct measures of body size (based on central coordinates; n=44), **(B)** number of assemblage time-series in each study, **(C)** species richness observed across assemblages, **(D)** total number of body size observations across assemblages, **(E)** duration of sampling, and taxonomic distribution of: **(F)** species represented, **(G)** body size observations and **(H)** assemblage time-series.

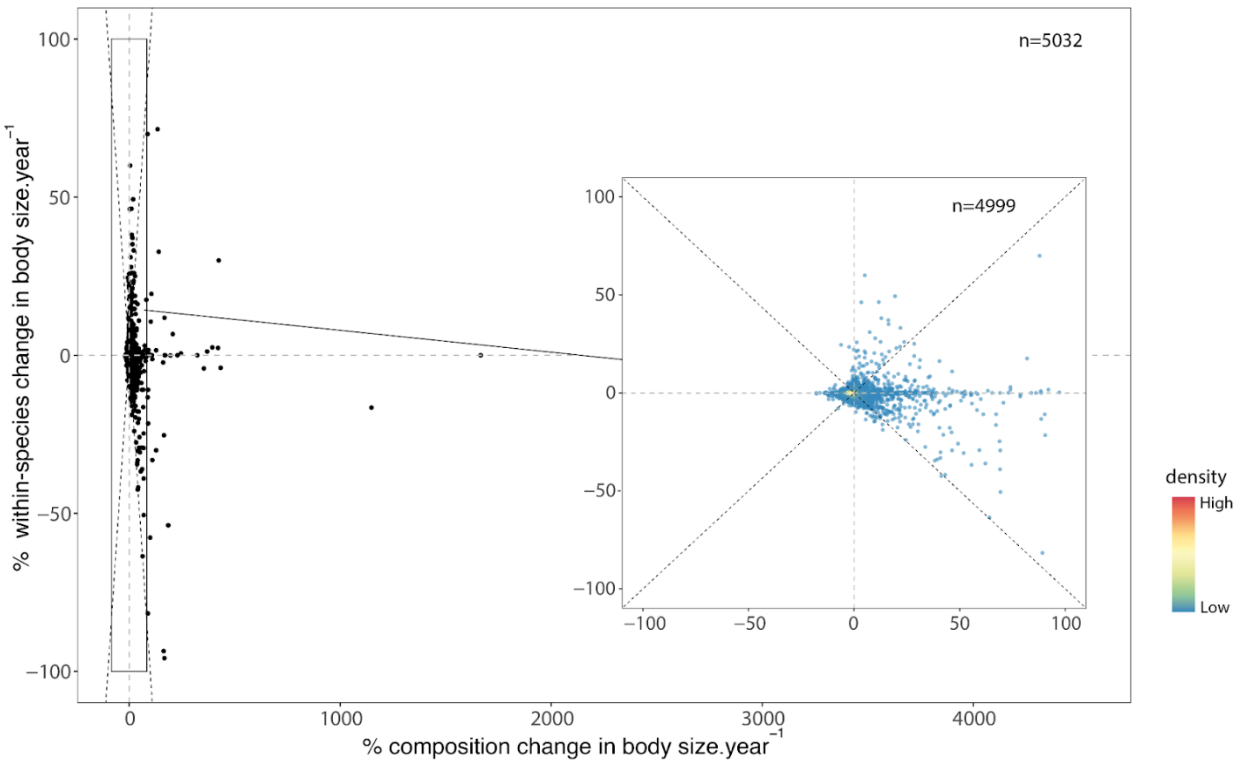


**Fig. S2.**

**Patterns of body size change through time in 5,032 assemblages.** Relationship between population-level changes and community-level changes. Both axes show % changes standardised by the number of years between the first and last year sample on the assemblage (duration); assemblages (points) are coloured by density break (colder colours indicating lower densities); Dashed lines show x = 0, y = 0, x = y and y=-x. Please see fig. S5 for more details.

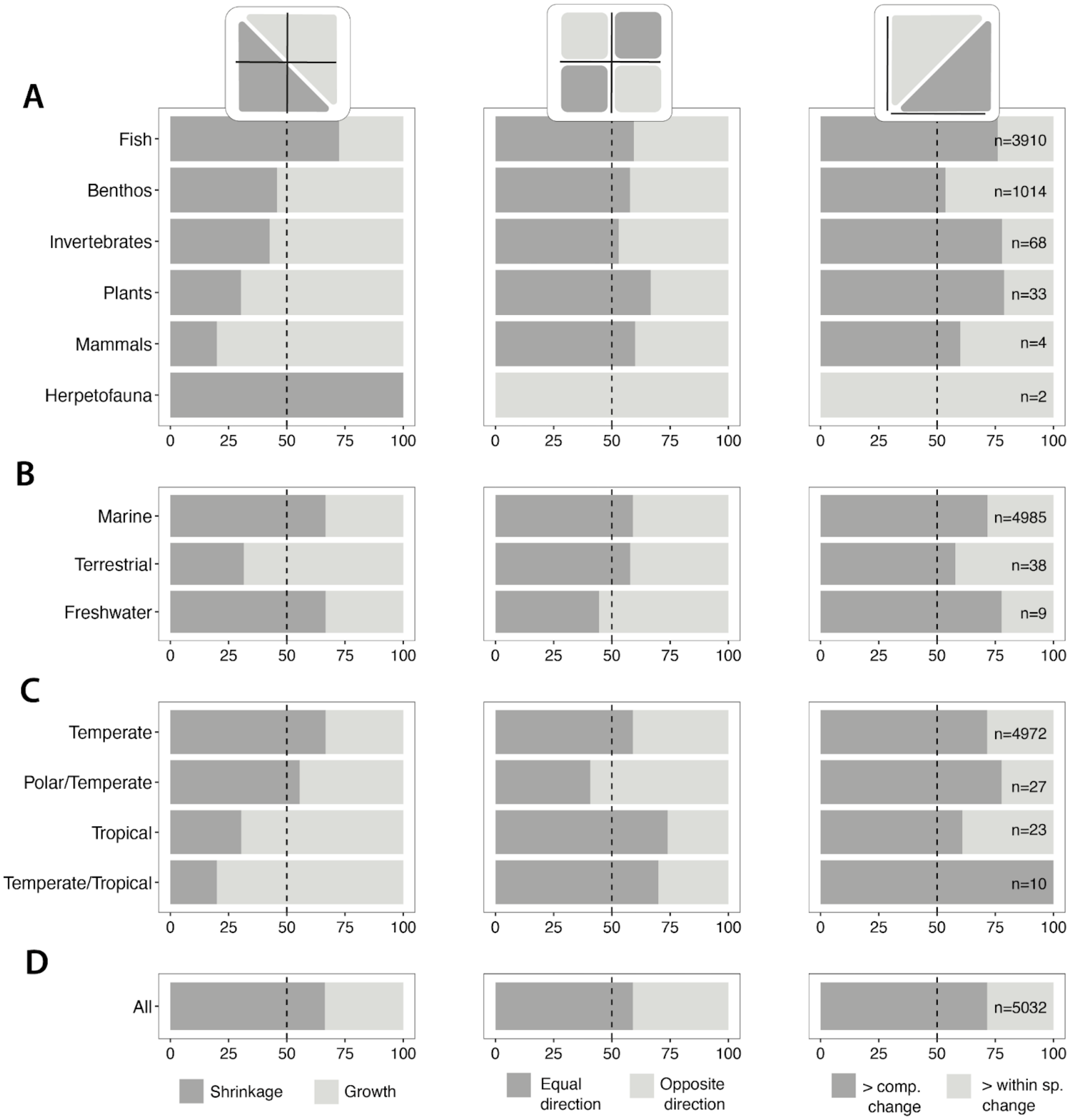


**Fig. S3.**

**Patterns of temporal body size change vary across (A) taxa, (B) realms, (C) climates, and the (D) globe.** Plots show the frequency distributions (in percentage) of the number of assemblages across different groups for each scenario depicted in Fig. 2B. Dashed lines mark the 50% threshold.

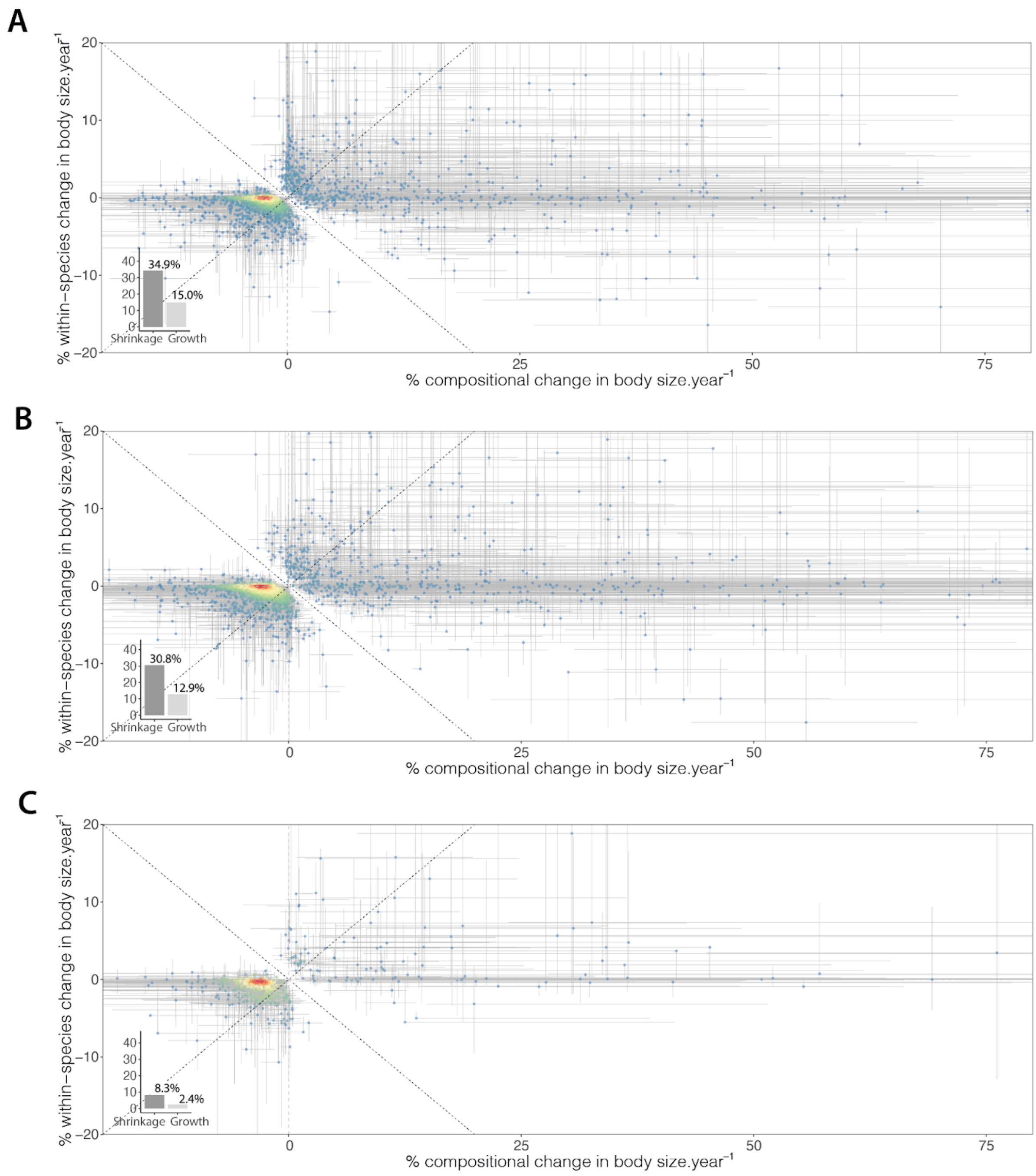


**Fig. S4.**

**Patterns of body size shrinkage through time are not influenced by start and/or end points.** Results using alternative start and/or end points. Analysis in the main text only compared the first and the last year of the time-series. The sensitivity analyses used instead an **(A)** fixed start year and random last year, a **(B)** random first year but fixed end year, or a **(C)** random start and end year. Points (assemblages) and grey lines indicate the assemblage median body size change and its IQR across 100 iterations (for each iteration changes were calculated using years chosen randomly according to the scenario assumptions). Points are coloured by density break (colder colours indicating lower densities). Inset histograms show % of assemblage where the median and IQR interval falls below (shrinkage) or above (growth) the y=-x line. Assemblages where the variation (IQR) crosses the y=-x line (and hence neither shrinking nor growing) are not shown.

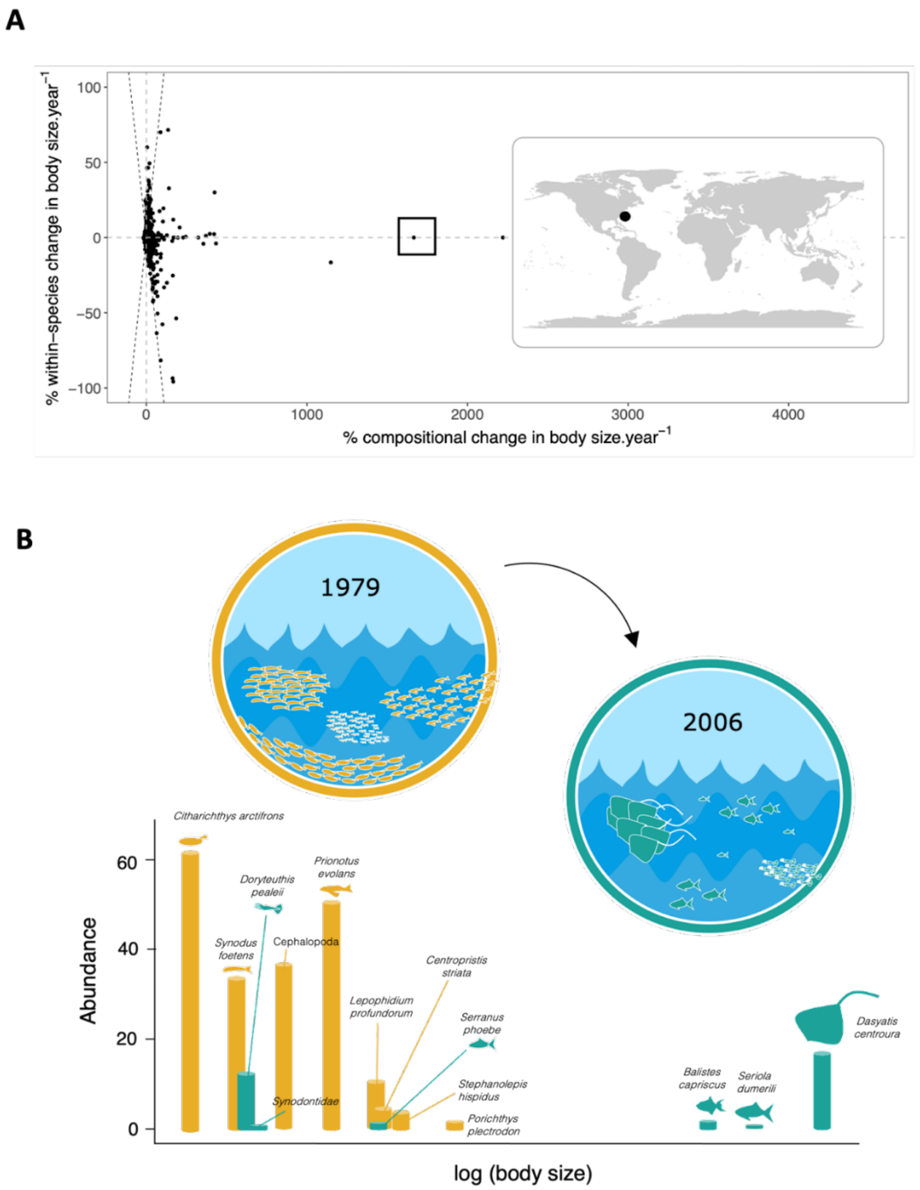


**Fig. S5.**

**Body size change through time: a look into the extremes.** The size of the assemblage through time will be determined by changes in species composition and within-species body size changes of each species. More extreme changes can occur in locations where assemblages are composed of species with very different body sizes to begin with, or/and when turnover occurs. For example, in the assemblage highlighted in **(A)** a wide spectrum of benthic organisms was sampled together. **(B)** In the first year (1979; orange), only small benthic organisms were recorded, although in high abundances ($CWM_{before}$=6.8x$10^{-3}$), however, by the last year (2006; green) a complete turnover in the assemblage had occurred. Despite decreases in both species' richness and abundance, the size of the assemblage increased ($CWM_{after}$=3.193), with the addition of several large-bodied species, including several individuals of the genus Dasyatis. This led to an assemblage ~467.5 times bigger (in weight) than the original, an increase of ~1666% per year due to compositional changes alone. Note that the illustrations are not to scale, and thus the size of the icons does not represent the real difference between the average body size of the species in the wild.

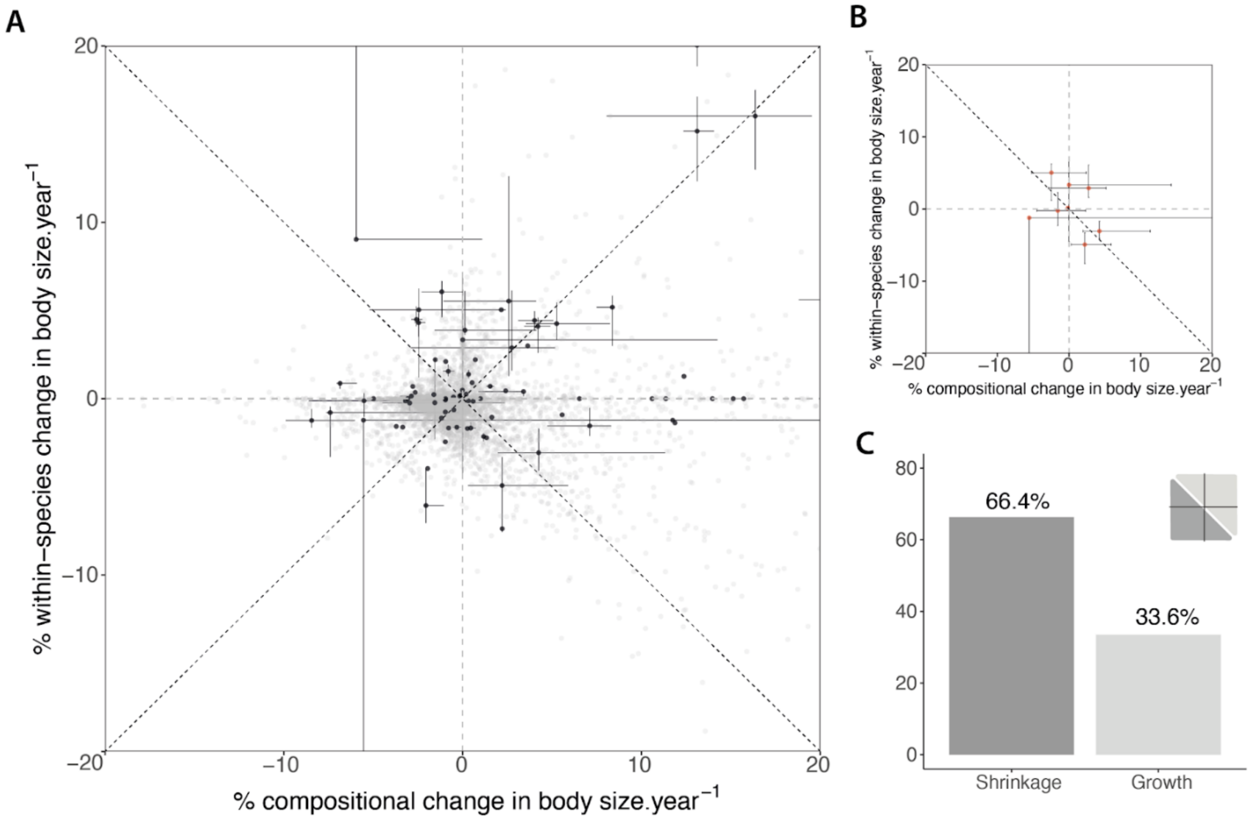


**Fig. S6.**
**Sensitivity analysis using different rarefaction subsets. (A)** only 1.77% of assemblages (black points; n=89) are affected by the sample-based rarefaction process. Grey lines indicate the variation (IQR) found in this subset of assemblages across 100 resamples. Assemblages unaffected by this process are shown in grey. **(B)** Assemblages where the variation (IQR) crosses the y=-x line (and hence neither shrinking nor growing; n=8). **(C)** Histograms show % of assemblage where the median and IQR interval falls below (shrinkage) or above (growth) the y=-x line after excluding assemblages highlighted in (**B**). For clarity, assemblages with % change.year$^{-1}$ higher than 20% are not shown in panels (A) and (B), but are included in (C).

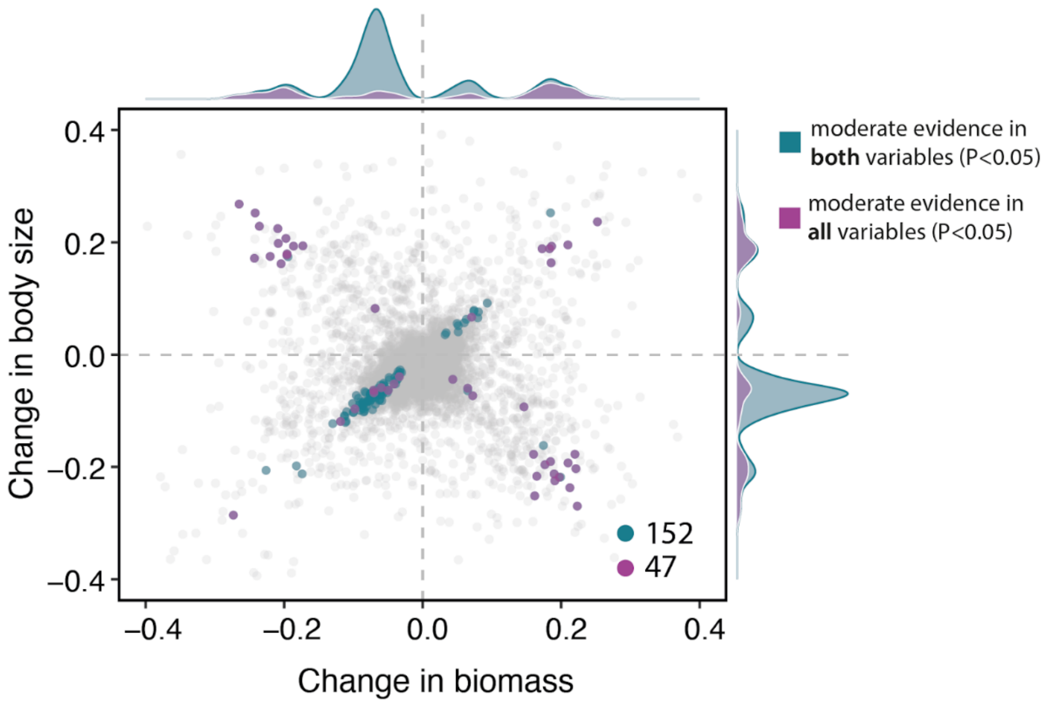


**Fig. S7.**

**Body size change through time as a function of biomass change.** Only assemblages for which strong or moderate evidence (P<0.05) was detected for both variables plotted are shown in blue. Purple highlights the assemblages for which significant changes through time were detected in all three variables (n=47), all remaining assemblages are shown in light grey.

### Supplementary Results

At large scales and often using species-level trait values, previous studies generally conclude that body size has, on average, been decreasing (*2, 3, 21*). This is due both to compositional changes, whereby bigger species are disproportionately replaced by smaller species (*3*) and within-species changes associated with the removal of larger individuals (*35*). However, data limitations on individual-level body size make it difficult to assess the importance (and signal) of the latter, thus, constraining global assessments to quantify body size change using species-level trait values alone. Here, we follow a similar approach but use BioTIME data to test if the assemblage body size shrinkage patterns presented in the main text are observed at the global scale when using different types of trait data.

For this analysis, we used two distinct subsets of BioTIME data: the 44 studies with directly measured estimates of body size (as described in the Materials and Methods; hereafter called type 1 data), and a larger subset of studies for which estimates of body size could be retrieved from major published databases (species trait averages; hereafter called type 2 data). Note that when matching with trait databases (i.e., the type 2 data), we did not work with only the subset of 44 studies featured in the main text but considered instead all BioTIME data that reported counts of the number of individuals and met our duration criteria. The different steps of data preparation and analysis for the former (type 1) are summarised in the Materials and Methods (*17*), any additional data, statistical analyses and supplementary results are described in detail below.

Additional Trait data (Trait databases)

Five open-access global databases were identified as having partially overlapping observations with species listed in the BioTime dataset: AmphiBIO (*44*); https://doi.org/10.6084/m9.figshare.4644424.v5), TRY database (*45*); https://www.try-db.org/), EltonTraits 1.0 (*46*); https://doi.org/10.6084/m9.figshare.c.3306933.v1), FishBase (*47*); www.fishbase.org) and Carabids.org (*48*); http://www.carabids.org). While a number of traits related to body size are available in these datasets, here, and given the broad-scale goal of the paper, we choose to select the body size trait that had the higher cover for each taxon. For birds and mammals, it was body mass ('*body_size_mm'* field). However, for ectotherms, length was used instead, as it is a more commonly available measure and considered more reliable than body mass. For fish and beetles (hereafter: Invertebrates) we used maximum body length (*'MaxLengthTL'* and *'maxSize'* fields, respectively), for amphibians we used snout-vent length ('*body_size_mm'* field), and for plants we used plant height (*'veg_height'* field). Only species-level trait values were considered and when there were multiple values for a particular species (i.e., TRY database), we took the median.

Synthesis and harmonisation of data

To merge and harmonise the assemblage time-series and trait databases data and optimise species matching, we first followed a series of steps to deal with species names and incompatibilities between the two sources of data. As a preliminary step, we reviewed species names listed in BioTIME with the taxize R package (*49*). An additional field was created with potential synonyms, or alternative scientific names (identified misspelling errors). Finally, common names were also flagged and, when possible, converted to scientific synonyms using additional data sources (ITIS and NCBI), helped by manual inspection based on the description provided. This process allowed for 913 species (11%) extra matches between the two sources of data. Because the sample-based rarefaction could result in different species composition for some assemblage time-series, we decided to work with all 100 BioTIME resamples (see '*Sensitivity analysis*' section in Material and Methods (*17*)). The matching was then done for each BioTIME resample dataset separately, and filtered to keep only records from where trait values were available (i.e., common species across datasets). On average, we retained records for ~5,400 species across ~21,000 assemblage time-series (fig. S8), with an average of ~72% completeness (i.e., proportion of species in the assemblage time-series from which trait data was retrieved).

*Calculating body size*

For assemblages time-series matched with trait databases data (type 2 data), average body size of individuals in a given year ($\widehat{BS}_{r,t}$) was calculated, by averaging all species' individual body sizes, weighted by their abundances within the assemblage. For details on how $\widehat{BS}_{r,t}$ was calculated for the 44 studies with directly measured estimates of body size (type 1 data) see '*Body size, abundance, and biomass change*' section in Materials and Methods (*17*). To make all body size estimates comparable, all values were standardised using classic z-scores, where individual observations in a group are scaled relative to the mean and standard deviation of all observations of that group. This was done for each assemblage time-series separately.

Statistical analyses - Models of body size change

We explored the global patterns of body size change using mixed-effects models. Year (mean-centered) was included as a fixed effect, and was also included as a random slope varying across studies and assemblages, with assemblages nested into the original studies from which they originated in order to account for the non-independence of the time-series. All statistical models were fitted in a Bayesian framework using the package 'brms' (*50*) in R (v3.6.3; (*43*)). We modelled average individual body size change assuming a skew-normal distribution and an identity link function. The overall model structure implemented using the bms syntax was:

$$y_{j,i,t} \sim SkewNormal(\mu_{j,i,t}, \sigma, a),$$

$$\mu_{j,i,t} = \beta_0 + \beta_{0j} + \beta_{0j,i} + (\beta_1 + \beta_{1j} + \beta_{1j,i})year_{j,i,t},$$

where $y_{j,i,t}$ is the average individual body size change in year $t$ of the $i$th assemblage in the $j$th study. $year_{j,i,t}$ is the time in years, $\beta_0$ and $\beta_1$ are the global intercept and slope (fixed effects), $\beta_{0j}$and $\beta_{1j}$ are the study-level departures from $\beta_0$ and $\beta_1$ (respectively; study-level random intercept and slope), and $\beta_{0j,i}$ and $\beta_{1j,i}$ are the (nested) assemblage-level departures from $\beta_{0j,i}$ and $\beta_{1j,i}$ (respectively; assemblage-level random intercept and slope). We used weakly regularizing priors for the global intercept and slope, residual variation ($\sigma$), and skew parameter ($a$):

$$\beta_0 \sim N(0,2),$$

$$\beta_1 \sim N(0,1),$$

$$\sigma \sim student\ t(3, 0, 2.5),$$

$$a \sim N(0, 4).$$

Group level parameters were drawn from the student-t distribution:

$$\sigma_{0j} = \sigma_{1j} = \sigma_{0ji} = \sigma_{1ji} \sim student\ t(3, 0, 2.5).$$

Correlations between levels of the grouping-factors were estimated using the Cholesky decomposition (L) of the correlation matrix, with a Lewandowski-Dorota- Joe (LKJ) prior:

$$L \sim LKJ(2).$$

The model was fit to 100 resamples of each dataset (i.e. type 1 and type 2) to adjust for any variation in species composition arising when sample effort was standardised using sample-based rarefaction. For each model fit, we extracted the 100 draws from the posterior distribution, which were combined for making inferences.

Results

In this analysis, we found no evidence of systematic body size change across taxa, realms and the globe (fig. S9 and S10). Only a few studies (mostly of marine fish) show some departure from zero. We found similar results independent of the type of body size trait data, i.e., we cannot detect change either when we use species mean body sizes from trait databases (type 2 data, 20,925 assemblages, fig S1) or when we use field body size measurements reported in BioTIME (type 1 data, 5,032 assemblages, fig S8).

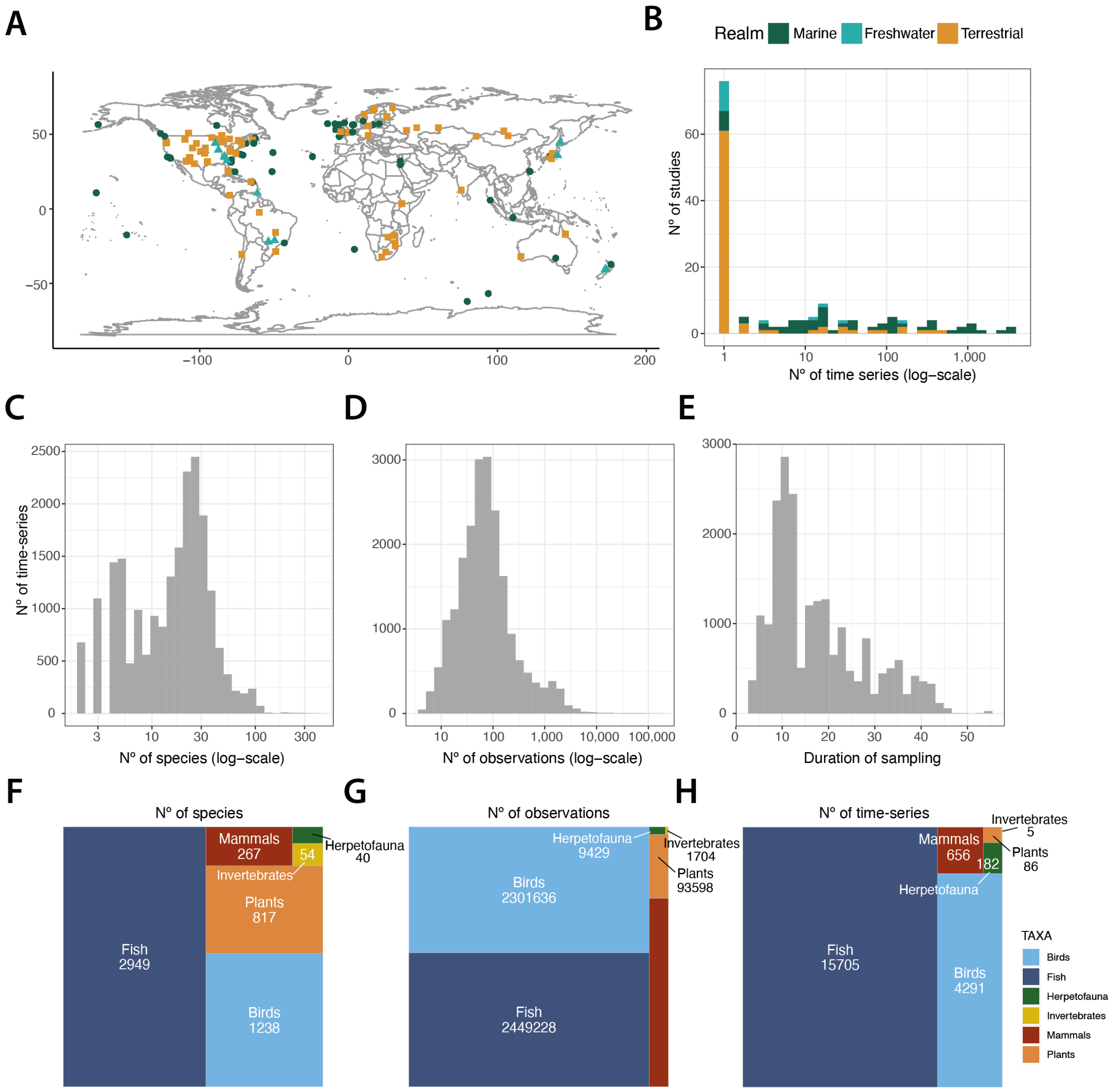


**Fig. S8.**

**Distribution of study data characteristics (BioTIME data, type 2). (A)** location of the studies with indirect measures of body size (based on central coordinates; n=151), **(B)** number of assemblage time-series in each study, **(C)** species richness observed across assemblages, **(D)** total number of body size observations across assemblages, **(E)** duration of sampling, and taxonomic distribution of: **(F)** species represented, **(G)** body size observations and **(H)** biodiversity time-series.

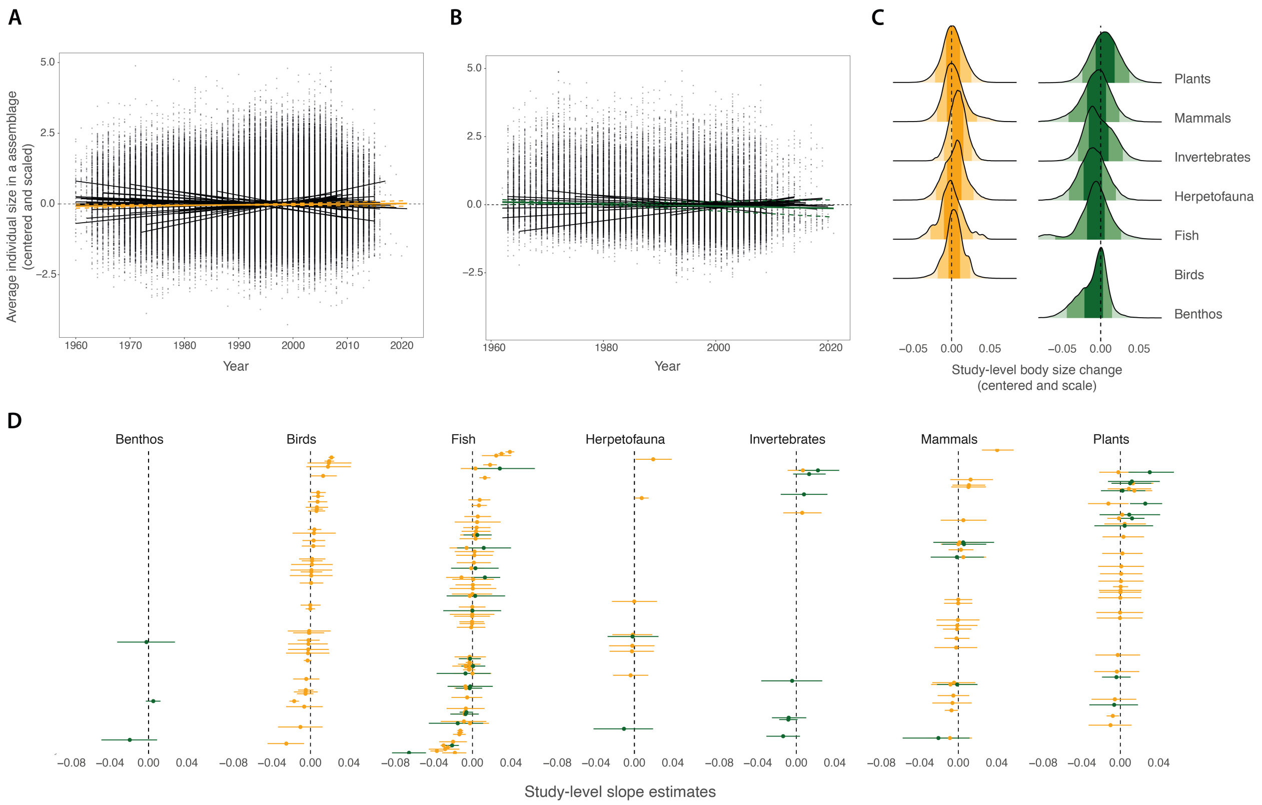


**Fig. S9.**

**No clear global temporal trend in body size change across assemblages.** Overall trend in average individual body size (centred and scaled) change as a function of year when considering **(A)** species' average body size estimates or **(B)** direct measurements of size. Lines depict the global median and 90% credible interval across all assemblages. Black lines show study-level variation. **(C)** Density ridges of posterior distributions of the study-level slope coefficients for a given taxon. **(D)** Estimates of change for each taxon, each point represents a single study, with the bar showing the 90% credible interval; studies are arranged by their median value (point). In all plots, colour represents the type of body size data: orange = average body size from trait databases (20,925 assemblages across 151 studies), dark green = body size from direct measurements as reported in the BioTIME database (5,032 assemblages across 44 studies).

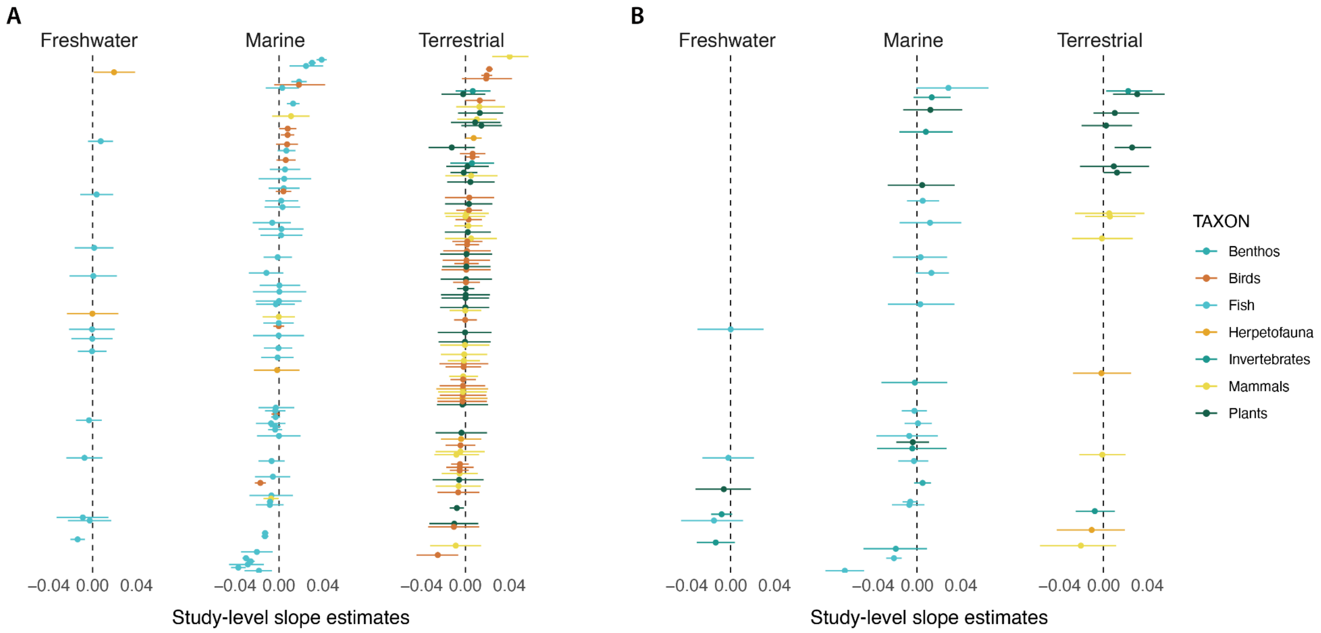


**Fig. S10.**

**Patterns of average individual body size change across the different realms.** Estimates of change for each realm when considering **(A)** species' average body size estimates (151 studies) or **(B)** direct measurements of size (44 studies). Each data point represents a single study, with the bar showing the 90% credible interval; studies are arranged by their median change value (point).

**Table S1.** Details on the datasets used in this study.

| Study ID | Start year | End year | Nr yrs | Taxon | Realm | Climate | type1 | type2 | References |
|---|---|---|---|---|---|---|---|---|---|
| 39 | 1970 | 2015 | 45 | Birds | Terrestrial | Temperate | | x | (*51-54*) |
| 42 | 1960 | 1979 | 20 | Birds | Terrestrial | Temperate | | x | (*55-61*) |
| 44 | 1962 | 1977 | 16 | Plants | Terrestrial | Temperate | x | x | (*62-64*) |
| 45 | 2006 | 2019 | 14 | Fish | Marine | Tropical | x | x | (*65, 66*) |
| 46 | 1960 | 1979 | 20 | Birds | Terrestrial | Temperate | | x | (*67*) |
| 47 | 1960 | 1977 | 18 | Birds | Terrestrial | Temperate | | x | (*68*) |
| 51 | 1964 | 1977 | 14 | Birds | Terrestrial | Temperate | | x | (*69, 70*) |
| 52 | 1968 | 1980 | 13 | Mammals | Terrestrial | Polar | | x | (*71, 72*) |
| 53 | 1966 | 1976 | 10 | Mammals | Terrestrial | Temperate | | x | (*73, 74*) |
| 56 | 1989 | 2019 | 31 | Mammals | Terrestrial | Temperate | x | x | (*75*) |
| 57 | 1981 | 2012 | 32 | Fish | Freshwater | Temperate | | x | (*76*) |
| 58 | 1991 | 2008 | 18 | Birds | Terrestrial | Tropical | | x | (*77, 78*) |
| 59 | 1977 | 2002 | 26 | Mammals | Terrestrial | Temperate | | x | (*79*) |
| 60 | 1982 | 2005 | 6 | Plants | Terrestrial | Tropical | | x | (*80-85*) |
| 67 | 1983 | 2006 | 24 | Birds | Terrestrial | Temperate | | x | (*86*) |
| 81 | 1997 | 2010 | 14 | Mammals | Marine | Temperate | | x | (*87, 88*) |
| 86 | 1985 | 2007 | 16 | Plants | Marine | Temperate | x | | (*89*) |
| 91 | 1992 | 1999 | 8 | Birds | Marine | Temperate | | x | (*90*) |
| 100 | 1981 | 2011 | 31 | Fish | Marine | Temperate | | x | (*91-93*) |
| 108 | 1981 | 2006 | 25 | Birds | Marine | Global | | x | (*94*) |
| 112 | 1973 | 2005 | 22 | Fish | Marine | Temperate/Tropical | | x | (*95*) |
| 119 | 1970 | 2010 | 41 | Fish | Marine | Temperate | x | x | (*96*) |
| 121 | 2000 | 2010 | 8 | Fish | Marine | Temperate/Tropical | | x | (*97*) |
| 123 | 2000 | 2009 | 10 | Fish | Marine | Temperate | x | x | (*98*) |
| 125 | 1988 | 2000 | 12 | Fish | Marine | Temperate | x | x | (*99*) |
| 126 | 1974 | 1980 | 7 | Fish | Marine | Temperate | | x | (*100*) |
| 127 | 1980 | 1989 | 10 | Fish | Marine | Temperate | x | x | (*101*). |
| ††163 | 1993 | 2004 | 12 | Benthos | Marine | Temperate | x | | |
| ††163 | 1993 | 2004 | 12 | Birds | Marine | Temperate | | x | (*102*) |
| ††163 | 1993 | 2004 | 12 | Fish | Marine | Temperate | | x | |
| †166 | 1966 | 1990 | 23 | Birds | Marine | Global | | x | (*103-108*) |
| †166 | 1970 | 1990 | 15 | Mammals | Marine | Global | | x | |
| †169 | 1987 | 2006 | 20 | Birds | Marine | Temperate | | x | |
| †169 | 1987 | 2003 | 7 | Fish | Marine | Temperate | | x | (*109-112*) |
| †169 | 1987 | 2006 | 20 | Mammals | Marine | Temperate | | x | |
| 171 | 1992 | 2008 | 17 | Mammals | Marine | Temperate/Tropical | | x | (*113*) |
| †172 | 2000 | 2009 | 10 | Birds | Marine | Temperate | | x | (*114-117*) |
| †172 | 1998 | 2009 | 12 | Mammals | Marine | Temperate | | x | |
| 176 | 1999 | 2010 | 12 | Invertebrates | Marine | Temperate | x | | (*118*) |
| 180 | 1970 | 1995 | 26 | Fish | Marine | Polar/Temperate | | x | (*119*) |
| 182 | 1988 | 2009 | 22 | Fish | Marine | Temperate | | x | (*120*) |
| 189 | 2001 | 2010 | 10 | Fish | Marine | Tropical | | x | (*121*) |
| 190 | 2001 | 2010 | 10 | Fish | Marine | Tropical | | x | (*122*) |
| 192 | 1960 | 1980 | 19 | Mammals | Marine | Polar/Temperate | | x | (*123*) |
| 195 | 1978 | 2007 | 30 | Birds | Terrestrial | Temperate | | x | (*124*) |
| 195 | 1978 | 2007 | 30 | Plants | Terrestrial | Temperate | | x | |
| 196 | 1987 | 2012 | 12 | Fish | Marine | Temperate | | x | (*125*) |
| 197 | 1985 | 2013 | 28 | Fish | Marine | Temperate | | x | (*126*) |
| 198 | 1991 | 2013 | 23 | Fish | Marine | Temperate | | x | (*127*) |

| | | | | | | | | | |
|---|---|---|---|---|---|---|---|---|---|
| 205 | 2001 | 2009 | 8 | Fish | Marine | Temperate | | x | (*128*) |
| 206 | 1993 | 2008 | 16 | Fish | Marine | Temperate | | x | (*129*) |
| 207 | 2003 | 2008 | 6 | Fish | Marine | Temperate | | x | (*130*) |
| 208 | 1997 | 2007 | 11 | Fish | Marine | Temperate | | x | (*131*) |
| 209 | 1987 | 2010 | 24 | Fish | Marine | Temperate | | x | (*132*) |
| 210 | 1965 | 2011 | 47 | Fish | Marine | Temperate | | x | (*133*) |
| 211 | 1980 | 1987 | 8 | Fish | Marine | Temperate | x | x | (*134*) |
| 212 | 1973 | 1980 | 8 | Fish | Marine | Temperate | x | x | (*135*) |
| 213 | 1963 | 2008 | 46 | Fish | Marine | Temperate | x | x | (*136*) |
| 214 | 1960 | 2010 | 46 | Plants | Terrestrial | Temperate | | x | (*137*) |
| 215 | 1960 | 2008 | 49 | Birds | Terrestrial | Temperate/Tropical | | x | (*138*) |
| 216 | 2001 | 2005 | 5 | Birds | Terrestrial | Temperate | | x | (*139*) |
| 217 | 1992 | 2006 | 14 | Birds | Terrestrial | Temperate | | x | (*140*) |
| 218 | 2006 | 2010 | 5 | Birds | Terrestrial | Temperate | | x | (*141*) |
| 219 | 1995 | 2011 | 17 | Herpetofauna | Terrestrial | Temperate | | x | (*142*) |
| 220 | 1995 | 2011 | 17 | Birds | Terrestrial | Temperate | | x | (*142*) |
| 229 | 1988 | 2013 | 26 | Fish | Freshwater | Temperate | | x | (*143*) |
| †231 | 2000 | 2005 | 6 | Fish | Marine | Temperate | | x | |
| †231 | 2000 | 2005 | 6 | Herpetofauna | Marine | Temperate | | x | (*144*) |
| †231 | 2000 | 2005 | 6 | Plants | Marine | Temperate | | x | |
| 232 | 1968 | 1999 | 25 | Fish | Marine | Polar/Temperate | x | | (*145*) |
| 234 | 1965 | 2002 | 7 | Plants | Terrestrial | Temperate | x | x | (*146-152*) |
| 236 | 1995 | 2006 | 12 | Fish | Freshwater | Temperate | | x | (*153*) |
| 240 | 2003 | 2015 | 13 | Plants | Terrestrial | Temperate | x | x | (*154*) |
| 242 | 1998 | 2006 | 7 | Plants | Terrestrial | Temperate | | x | (*155*) |
| 243 | 1992 | 2014 | 22 | Plants | Terrestrial | Temperate | x | x | (*156, 157*) |
| 244 | 1999 | 2012 | 14 | Birds | Marine | Temperate | | x | (*158*) |
| 246 | 2000 | 2014 | 15 | Fish | Marine | Temperate | | x | (*159*) |
| 247 | 1975 | 2020 | 46 | Invertebrates | Freshwater | Temperate | x | | (*160*) |
| 248 | 2000 | 2012 | 13 | Plants | Terrestrial | Temperate | | x | (*161*) |
| 249 | 1992 | 2015 | 24 | Invertebrates | Terrestrial | Temperate | | x | (*162*) |
| 252 | 1978 | 1989 | 12 | Fish | Marine | Temperate | x | x | (*163*) |
| 254 | 1995 | 2018 | 24 | Plants | Freshwater | Temperate | x | | (*164*) |
| 255 | 1989 | 2007 | 10 | Plants | Terrestrial | Temperate | | x | (*165*) |
| 256 | 1987 | 2010 | 24 | Fish | Marine | Temperate | | x | (*132*) |
| 271 | 2000 | 2014 | 15 | Fish | Marine | Temperate | | x | (*166*) |
| 275 | 2009 | 2014 | 6 | Herpetofauna | Terrestrial | Temperate | x | | (*167*). |
| 277 | 1979 | 2008 | 10 | Plants | Terrestrial | Temperate | | x | (*62-64*) |
| 279 | 1979 | 2008 | 10 | Plants | Terrestrial | Temperate | x | x | (*62, 63, 168*) |
| 288 | 1970 | 2005 | 29 | Fish | Marine | Temperate | | x | (*96*) |
| 295 | 2008 | 2016 | 9 | Fish | Marine | Temperate | x | x | (*169*) |
| 296 | 2008 | 2016 | 9 | Fish | Marine | Temperate | | x | (*169*) |
| 305 | 1976 | 1990 | 15 | Herpetofauna | Terrestrial | Temperate | | x | (*170*) |
| 308 | 1980 | 1998 | 19 | Mammals | Terrestrial | Temperate | x | x | (*171*) |
| 311 | 1981 | 2013 | 33 | Mammals | Terrestrial | Temperate | | x | (*172*) |
| 312 | 1961 | 1984 | 8 | Mammals | Terrestrial | Tropical | | x | (*173*) |
| 316 | 1989 | 2006 | 18 | Herpetofauna | Terrestrial | Temperate | x | | (*174*) |
| 317 | 1995 | 2005 | 9 | Plants | Terrestrial | Temperate | | x | (*175*) |
| †318 | 1994 | 2009 | 13 | Birds | Terrestrial | Temperate | | x | (*176*) |
| †318 | 1994 | 2009 | 13 | Mammals | Terrestrial | Temperate | | x | |
| 319 | 1990 | 2003 | 13 | Herpetofauna | Terrestrial | Temperate | | x | (*177*) |
| 321 | 1995 | 2007 | 13 | Mammals | Terrestrial | Temperate | x | x | (*178*). |

| | | | | | | | | | |
|---|---|---|---|---|---|---|---|---|---|
| 324 | 2003 | 2007 | 5 | Plants | Terrestrial | Tropical | | x | (*179*) |
| 325 | 2003 | 2007 | 5 | Plants | Terrestrial | Tropical | | x | (*179*) |
| 327 | 1989 | 2005 | 17 | Mammals | Terrestrial | Temperate | x | x | (*180*) |
| 328 | 1979 | 2008 | 30 | Herpetofauna | Freshwater | Temperate | | x | (*181*) |
| 329 | 1990 | 2010 | 6 | Plants | Terrestrial | Tropical | | x | (*181*) |
| 332 | 1984 | 1995 | 12 | Fish | Freshwater | Temperate | | x | (*182*) |
| 335 | 1962 | 1974 | 12 | Fish | Freshwater | Temperate | | x | (*183*) |
| 336 | 1989 | 2002 | 14 | Plants | Terrestrial | Temperate | | x | (*79*) |
| 339 | 1960 | 2009 | 50 | Birds | Terrestrial | Temperate | | x | (*184*) |
| 340 | 1995 | 2009 | 15 | Plants | Terrestrial | Temperate | | x | (*185*) |
| 348 | 2006 | 2016 | 10 | Mammals | Terrestrial | Temperate/Tropical | x | x | (*186*) |
| 350 | 1989 | 2011 | 12 | Benthos | Marine | Temperate | x | | (*187, 188*) |
| 351 | 1989 | 2004 | 13 | Benthos | Marine | Temperate | x | | (*189, 190*) |
| 356 | 1971 | 2013 | 34 | Plants | Terrestrial | Tropical | | x | (*191*) |
| 357 | 1994 | 2006 | 13 | Mammals | Terrestrial | Temperate | | x | (*192*) |
| 358 | 1977 | 1992 | 16 | Birds | Terrestrial | Temperate | | x | (*193*) |
| 359 | 2000 | 2012 | 13 | Fish | Marine | Temperate | | x | (*194*) |
| 361 | 1962 | 1983 | 22 | Birds | Terrestrial | Temperate | | x | (*195*) |
| 363 | 1963 | 1999 | 37 | Birds | Terrestrial | Temperate | | x | (*196*) |
| 365 | 1997 | 2002 | 6 | Fish | Marine | Temperate | | x | (*197*) |
| 366 | 1989 | 2013 | 25 | Mammals | Terrestrial | Temperate | | x | (*198*) |
| 372 | 2005 | 2013 | 9 | Birds | Terrestrial | Temperate | | x | (*199*). |
| 373 | 2005 | 2012 | 8 | Mammals | Terrestrial | Temperate | | x | (*200*) |
| 374 | 2004 | 2014 | 11 | Birds | Marine | Temperate | | x | (*201*) |
| 375 | 2004 | 2014 | 11 | Invertebrates | Terrestrial | Temperate | x | x | (*202*) |
| 377 | 2009 | 2013 | 5 | Birds | Terrestrial | Temperate | | x | (*202*) |
| 381 | 1986 | 1992 | 7 | Herpetofauna | Terrestrial | Temperate | | x | (*203*) |
| 382 | 1960 | 1967 | 8 | Mammals | Terrestrial | Temperate | | x | (*204, 205*) |
| 402 | 2010 | 2015 | 6 | Fish | Freshwater | Tropical | | x | (*206*) |
| 403 | 1989 | 1995 | 7 | Herpetofauna | Freshwater | Tropical | | x | (*207*). |
| 412 | 1995 | 2000 | 6 | Fish | Marine | Temperate | | x | (*208*) |
| 420 | 1964 | 2001 | 38 | Birds | Terrestrial | Polar/Temperate | | x | (*209*) |
| 427 | 1977 | 2008 | 32 | Invertebrates | Freshwater | Temperate | x | | (*210, 211*) |
| †428 | 1963 | 2015 | 45 | Birds | Marine | Temperate | | x | (*212-215*) |
| †428 | 1960 | 2015 | 56 | Fish | Marine | Temperate | | x | |
| 430 | 1985 | 2016 | 32 | Fish | Freshwater | Temperate | | x | (*216*) |
| 431 | 1984 | 2015 | 30 | Fish | Freshwater | Temperate | | x | (*216*) |
| 432 | 1998 | 2016 | 19 | Fish | Freshwater | Temperate | | x | (*216*) |
| 435 | 2012 | 2019 | 7 | Fish | Marine | Polar/Temperate | | x | (*217*) |
| 435 | 2009 | 2019 | 11 | Invertebrates | Marine | Polar/Temperate | x | | |
| 436 | 2005 | 2012 | 5 | Fish | Marine | Tropical | x | x | (*218*) |
| 438 | 2006 | 2014 | 7 | Fish | Marine | Tropical | x | x | (*219*) |
| 439 | 1985 | 1997 | 13 | Birds | Terrestrial | Temperate | | x | (*220*) |
| 440 | 1985 | 1997 | 13 | Birds | Terrestrial | Temperate | | x | (*220*) |
| 441 | 1985 | 1997 | 13 | Birds | Terrestrial | Temperate | | x | (*220*) |
| 442 | 1980 | 1985 | 6 | Birds | Terrestrial | Temperate | | x | (*221*) |
| 444 | 1983 | 1987 | 5 | Birds | Terrestrial | Temperate | | x | (*222*) |
| 446 | 2007 | 2011 | 5 | Mammals | Terrestrial | Temperate | | x | (*223*) |
| 447 | 2006 | 2014 | 9 | Mammals | Terrestrial | Temperate | | x | (*224*) |
| 449 | 2000 | 2009 | 10 | Mammals | Terrestrial | Temperate | | x | (*225*) |
| 464 | 2007 | 2011 | 5 | Plants | Terrestrial | Temperate | | x | (*226-228*) |
| 465 | 2007 | 2012 | 6 | Plants | Terrestrial | Temperate | | x | (*226-228*) |

| | | | | | | | | | |
|---|---|---|---|---|---|---|---|---|---|
| 466 | 1986 | 2008 | 23 | Fish | Marine | Temperate | | x | (*229*) |
| 469 | 1985 | 2011 | 27 | Fish | Marine | Temperate | | x | (*230*) |
| 471 | 2004 | 2013 | 10 | Plants | Terrestrial | Temperate | | x | (*231*) |
| 475 | 1960 | 1972 | 12 | Birds | Terrestrial | Temperate | | x | (*232*) |
| 477 | 1998 | 2012 | 15 | Invertebrates | Marine | Temperate | x | | (*233, 234*) |
| 501 | 2004 | 2012 | 9 | Fish | Marine | Temperate | | x | (*235*) |
| 502 | 1962 | 2009 | 7 | Plants | Terrestrial | Temperate | x | x | (*236*) |
| 504 | 2003 | 2007 | 5 | Fish | Marine | Temperate | | x | (*237*) |
| 505 | 1990 | 2012 | 11 | Fish | Marine | Temperate | | x | (*238*) |
| 511 | 2005 | 2015 | 6 | Fish | Marine | Tropical | x | x | (*239*) |
| 516 | 1997 | 2013 | 5 | Mammals | Terrestrial | Tropical | | x | (*240-244*) |
| 521 | 1972 | 2017 | 37 | Mammals | Terrestrial | Temperate | | x | (*245*) |
| 522 | 1987 | 2013 | 20 | Birds | Terrestrial | Temperate | | x | (*246*) |
| 523 | 1992 | 2007 | 16 | Birds | Terrestrial | Temperate | | x | (*247*) |
| †524 | 1993 | 1998 | 6 | Birds | Terrestrial | Tropical | | x | (*248*) |
| †524 | 1993 | 1998 | 6 | Mammals | Terrestrial | Tropical | | x | |
| 525 | 2006 | 2013 | 7 | Fish | Marine | Temperate | | x | (*249*) |
| 526 | 2001 | 2015 | 11 | Birds | Marine | Polar/Temperate | | x | (*250*) |
| 527 | 2007 | 2015 | 9 | Mammals | Marine | Polar/Temperate | | x | (*251*) |
| 547 | 2007 | 2011 | 5 | Plants | Terrestrial | Temperate | | x | (*226-228*) |
| 548 | 2007 | 2012 | 6 | Plants | Terrestrial | Temperate | | x | (*226-228*) |
| 549 | 2011 | 2015 | 5 | Fish | Freshwater | Tropical | x | x | (*252, 253*) |
| 550 | 2005 | 2021 | 17 | Fish | Freshwater | Tropical | x | x | (*254*) |
| 551 | 1999 | 2018 | 19 | Fish | Freshwater | Tropical | | x | (*255*) |
| 551 | 1999 | 2018 | 20 | Fish | Freshwater | Tropical | x | | |
| 552 | 1963 | 2010 | 7 | Invertebrates | Terrestrial | Temperate | x | | (*256, 257*) |
| 3541064 | 2003 | 2015 | 13 | Plants | Marine | Temperate/Tropical | x | | (*258*) |
| 3541065 | 2008 | 2014 | 7 | Plants | Marine | Temperate/Tropical | x | | (*258*) |

Note: † These studies were classified as ‘multiple taxa’ in the original data sources; †† These studies were classified as ‘benthos’ in the original data sources.
